# Cheap, Robust and Versatile Two-Dimensional Chromatography system for Proteomics of Nanogram Scale Samples

**DOI:** 10.1101/2024.04.02.587751

**Authors:** Eduardo S. Kitano, Gareth Nisbet, Yana Demyanenko, Katarzyna M Kowalczyk, Louisa Iselin, Stephen Cross, Alfredo Castello, Shabaz Mohammed

## Abstract

In this work we describe a low loss fractionation system comprised of a reconfigured Evosep One LC system for the first dimension and a repurposed 3D-printer as a fraction collector. The setup operates as a high-pH fractionation system capable of effectively working with nanogram scales of lysate digests. The 2D RP-RP system demonstrated superior proteome coverage over single-shot data-dependent acquisition (DDA) analysis using only 5 ng of human cell lysate digest with performance increasing with increasing amounts of material. We found that the fractionation system allowed over 70% signal recovery at the peptide level and, more importantly, we observed over 30% increase on protein level intensity which indicates the complexity reduction afforded by the system outweighs the sample losses endured. The application of data-independent acquisition (DIA) and wide window acquisition (WWA) to fractionated samples allowed more than 8,000 proteins to be identified from 50 ng of material. The utility of the 2D system was further investigated for phosphoproteomics (>21,000 phosphosites from 50 μg starting material) and pull-down type experiments and showed substantial improvements over single-shot experiments. We show that the 2D RP-RP system is highly versatile and powerful tool for many proteomics workflows.

## INTRODUCTION

Technological advancements in Liquid Chromatography (LC)^1-4^ and Mass Spectrometry (MS)^5-7^ have significantly improved the depth of proteome coverage in bottom-up proteomics. The combination of these two analytical tools on the analysis of peptides from cells or tissues has become the state-of-the art workflow in proteomics. New mass spectrometry configurations^8, 9^ combined with robust nanoLC^10, 11^ has led to considerable increase in proteome coverage and throughput. Nevertheless, deep and comprehensive proteome characterization by single-shot LC-MS still represents a challenge. Two-dimensional (2D) separation strategies play a pivotal role obtaining a more complete characterization of complex biological samples by reducing sample complexity.^12^

Several fractionation strategies have been used in combination with LC-MS/MS. Cation exchange^13-15^ and reversed-phase (RP)^16-19^ have been the most commonly used multidimensional separation modes, as reviewed by Yuan et al.^12^. High-pH has proven popular due to ease of set up and the use of concatenation to improve orthogonality with the second dimension low-pH RP nanoLC separation.^17^ Moreover, high and low-pH 2D separation (2D RP-RP) benefits from the higher resolving power, compatible solvents between the first and second dimensions, and no need of desalting steps prior to the second dimension.

Despite the ability to unravel complex biological mixtures, a key limitation of 2DLC methodologies lie in the sample losses accrued when transferring between dimensions. Nonspecific binding of peptides to silica, metal, and plastic surfaces significantly contributes to peptide loss, compromising sensitivity and quantitative accuracy in LC-MS analysis. These sample losses restrict the use of 2D separations to studies where abundant material is readily available. For studies involving samples with limited availability, such as clinical specimens, rare cell populations, or those enriched for specific proteins/peptides, achieving adequate analytical depth while minimizing sample losses presents a significant challenge. Sample preparation approaches through the use of detergents have tried to minimise losses.^19, 20^ Also, others have developed nano/microscale automated fractionator systems^18, 21^, in an attempt to minimise losses during the transfer step of the fractions to second dimension separation which can also include the use of low-binding containers, low surface area glass nanowell chips and MS-compatible additives in downstream sample handling steps.^19, 21, 22^

A major challenge of fractionation system is the need for bespoke and/or costly fractionation equipment. Furthermore, robustness of nanoLC is substantially worse than high flow systems and require constant monitoring for carryover and separation performance. In this work, we reconfigured an Evosep One LC system^10^ for a high-pH separation and coupled it to a fractionation system built from an inexpensive 3D printer that is highly versatile and is capable of fractionating low peptide amounts from cell lysates.

## EXPERIMENTAL SECTION

### Sample preparation and protein digestion

Expi293F cell (Gibco) lysate was submitted to in-solution LysC/trypsin digestion as previously described^15^, while HeLa cell lysate was submitted to on-bead digestion using solid-phase enhanced sample-preparation (SP3) method.^23^

### Phosphopeptide enrichment

Desalted peptide sample from the digestion of HeLa cell lysate was submitted to phosphopeptide Zr-IMAC enrichment protocol as previously described.^24^

### RNA Interactome Capture and sample preparation

RNA-binding proteins (RBPs) present in HEK293 cells were profiled via RNA interactome capture (RIC) approach.^25, 26^ After treating with benzonase, RIC samples were processed via SP3 clean-up method followed by on-bead digestion as described before. For details, see the Supporting Information.

### Off-line high-pH reversed-phase nano-fractionation system

Peptide samples were loaded on Evotip pure and submitted to high-pH chromatography using an Evosep One system (Evosep Biosystems). Separation was performed in an in-house packed C-18 column and analysed using the 30SPD method. Mobile phases A and B consisted of 10 mM of triethylammonium bicarbonate (TEAB, pH8), and 100% ACN, respectively. Forty fractions were collected and concatenated into 8 main fractions in a 96-well plate at 1-minute intervals using a 3D-printer-based fraction collector, fabricated from a Creality Ender 5 S1 printer and powered by a Raspberry pi computer (details on https://github.com/garethnisbet/Fraction-Collection-Unit and Supporting Information).

### Nano-Liquid Chromatography, mass spectrometry and data processing

Peptides were separated on Ultimate 3000 RSLCnano system (Thermo Fisher Scientific) using an in-house packed 50μm ID x 50cm L Reprosil-Gold C-18 (Dr. Maisch) analytical column at 100nL/min. Eluting peptides were electro sprayed into an Orbitrap Ascend Tribrid mass spectrometer (Thermo Fisher Scientific), using Data Dependent (DDA), Data Independent (DIA) or Wide Window Acquisition (WWA) modes.

LC-MS/MS data were analysed by four different platforms using the same human proteome database (Uniprot, Proteome ID: UP000005640, downloaded in August 2022, 79,759 sequences). DDA datasets were processed with FragPipe computational platform with MSFragger^27, 28^. We opted to use MaxQuant^29^ for processing the DDA raw data from the phosphopeptide-enriched samples. DIA experiments were analysed by DIA-NN software^30^ using library-free search, and WWA experiments were processed by INFERYS^31^ rescoring and CHIMERYS, implemented in Proteome Discoverer 3.0 software (Thermo Fisher Scientific). All experiments used standard parameters and filtered to 1% FDR protein and peptide level. The LC and MS method details are described in Supporting Information. Raw files and results from this study have been deposited to ProteomeXchange Consortium via the PRIDE^32^ partner repository with the data set identifier PXD051148. Subsequent analysis of data was performed in the Perseus environment^33^ and GraphpadPrism.

## RESULTS AND DISCUSSION

### Designing a robust high-pH nanoLC system

We hypothesized the Evosep One could be a robust platform for nanoLC fractionation since the trap column is single use and elution is only up to 40% acetonitrile, eliminating the transfer of the more hydrophobic junk on to the analytical column. These two features substantially improve robustness and has established the Evosep One as a formidable choice for high throughput proteomics^10, 34^ and will allow low intervention fractionation. To convert this system for fractionation we need to assess the system’s ability to operate at a basic pH as this is now considered optimal for the first dimension of a 2D RP-RP system. The separation system is centred on the use of a disposable trap column, where peptides are separated and stored in a loop by the gradients formed by four low-pressure pumps and then this preformed gradient is pushed towards the analytical column for separation by a single high-pressure pump.^10^ The need for preforming gradients requires careful control of flowrates and pressures and so the system only offers pre-set optimized methods. Changing pH will affect the relationships between viscosity, pressure, and flowrate. First, we evaluated the performance of the LC using an in-house constructed 150 μm ID C-18 column (matching commercial option) at standard pH and pH8. At pH8 the backpressures generated by the system, irrespective of the solvent composition, were similar to standard conditions indicating that the addition of TEAB to the mobile phase is not an issue. Using the 30 SPD method resulted in a median peak width of 47 seconds at baseline, across a 44-minute gradient (Figure S1). Repeatability was similar to low-pH, indicating that the analytical column is not affected by the mildly basic pH (Figure S2). When inspecting retention time of peptides using a scatter plot from human cell lysate digestion that was separated using high and low-pH we observed a very loose correlation, the expected trend established by others (Figure S3).^16, 35^ The peak width and gradient length will allow 40 1-minute fractions and allowing 5 concatenations and 8 highly discrete fractions potentially allowing an order of magnitude increase in peak capacity over a single dimension system. Similar 2D nanoLC RP-RP approaches using similar columns came to similar conclusions.^18, 19^

Nanolitre flowrate fraction collection systems require dedicated low sample loss instrumentation. We hypothesized that a cheap 3D printer which contains sufficient control in 3 dimensions is capable of being converted into a fraction collector and work in combination with the Evosep system (Figure 1A and Figure S4A). We dismantled the printer head in the 3D printer and developed a new controller that is powered by a Raspberry pi computer. The software is written in Python and uses PyGame for the GUI, Numpy of mathematical operations and PySerial for communication over USB. For the source code and detailed explanations, refer to the project’s GitHub repository: https://github.com/garethnisbet/Fraction-Collection-Unit. Custom 3D printed components were manufactured and adapted to the system for attaching the column and supporting the collection plate (Figure S4B-C). The system was designed to be flexible and able to automatically collect and concatenate multiple fractions into the 0.2 mL wells of a 96-well plate or 96-Evotip rack, depending on the LC system used for the second-dimension separation (Figure 4SC). Due to the low collection volumes (in our case each fraction was collected in 500 nL packages at 1-minute intervals), the effluent was collected by submerging the column outlet (~1 mm) into approximately 50 μL of preloaded LC buffer (Figure 1B and C).

**Figure 1.**
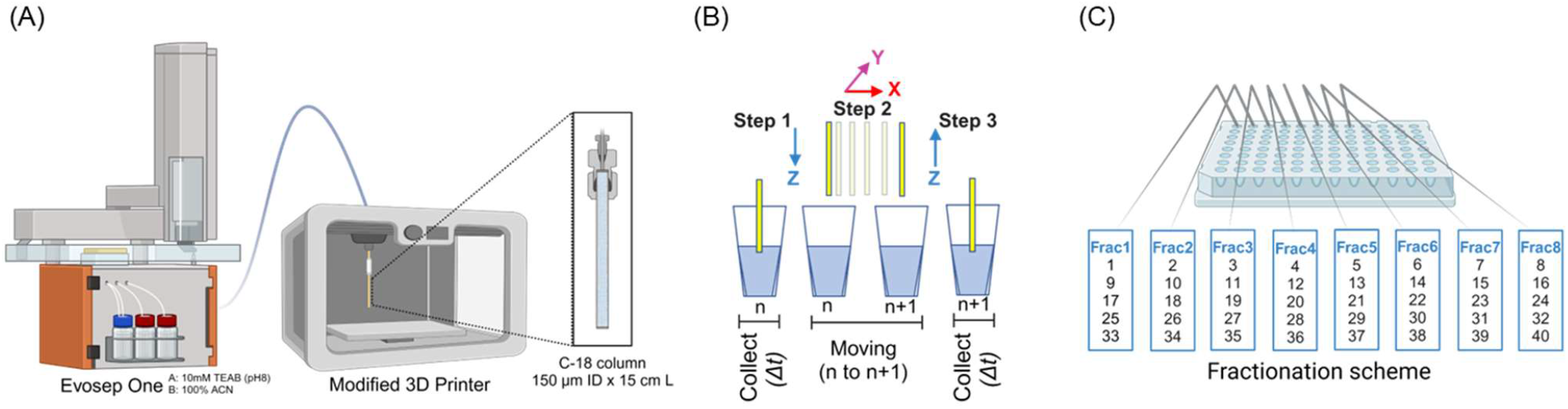
Off-line high-pH reversed-phase nano-fractionation system overview. (A) Peptide samples were submitted to first dimension separation using an Evosep One HPLC system equipped with an in-house packed 15cm L x 150 μm ID C-18 column coupled to a 3D-printer-based fraction collector. (B) XYZ moving mechanism of the fraction collector (position n to position n+1), showing the desired column outlet position. (C) Fractionation scheme showing the combination of 40 fractions, at 1-minute intervals, into eight concatenated main fractions (8x5 scheme). After first dimension separation, plates are inserted into the autosampler for the second dimension low-pH separation.

### Evaluating Fractionation approach

Having established a fractionation protocol, we then aimed to evaluate the performance and relative sensitivity of the 2D setup compared to a single-shot analysis. We opted to perform one ‘large scale’ (400 ng) fractionation and then use it to test a range of sample amounts varying from 0.25 ng to 200 ng of Expi293F digest. Table S1 lists the identified peptides categorized by sample amount. To maximize MS signal intensity and improve detection and selection of peaks for fragmentation, we employed different gradient times. For single-shot we employed gradients of 30 minutes for samples below 1 ng and 60 minutes for the rest. For fractionated samples we preferred 15 minutes for the analysis of peptide amounts below 1 ng (8 fractions, total 2h gradient time), and 30 minutes for the rest (total 4h). As expected, increasing sample amounts led to increased number of unique peptides and protein groups in both single-shot and 2D RP-RP analyses. Below 1 ng of sample, neither approach showed to be clearly superior although single-shot analyses potentially showed more favourable results (Figure 2A and B). In the lowest amount tested (0.25 ng), single-shot analysis resulted in 2,860 unique peptides and 663 protein groups, corresponding to 78% and 52% more IDs than obtained by 2D. Above and including 5 ng, the fractionation system produced more identifications. Initially, the 5-ng fractionation resulted in the identification of 23,908 unique peptides and 3,596 protein groups, representing increases of 10% and 8% compared to the single-shot analysis. Further increases in sample amount resulted in a progressive relative gain (Figure 2A) in identification of 2D over single-shot analysis. At 200 ng, 71,733 unique peptides and 7,231 proteins were identified, reflecting substantial improvements of 58% and 42% in comparison to single-shot. Important to note, single-shot identifications exhibited a tendency to plateau once past 25 ng of material while in 2D we did not observe such a levelling off for the amounts we applied suggesting we will continue seeing improved results with higher amounts of material. The Evosep loading tips are capable of handling up to 1 μg. It is noteworthy that 2D experiments produced higher peptide identifications at a lower average peptide intensity demonstrating the value of complexity reduction (Figure 2B). The fractionation approach resulted in a similar number of identified peptides across the fractions (Figure 2C), with most of peptides (70%) identified in one fraction (Figure 2D) which was similar to Kulak et al.^18^ and it confirms our choice of concatenation parameters. Comparing a single-shot analysis chromatogram to a collection of chromatograms for the associated fractionation sample show similar overall profile with the 2D experiment showing more peaks that are sharper indicating the superior separation power. 2D experiment resulted in an upper theoretical peak capacity value of 1,920, representing approximately 6 times improvement in comparison to our single-shot experiment and exceeding any single-shot analysis performed (Figure 2E).^36, 37^ Although we opted for a concatenation approach, the advent of ultra rapid mass analysers allow very short gradients (<5 minutes) and Guzman et al.^38^ have shown that non-concatenated fractionation can be orthogonal, powerful and simpler. Our fractionation system is compatible as we can change the fractionation collection pattern.

**Figure 2.**
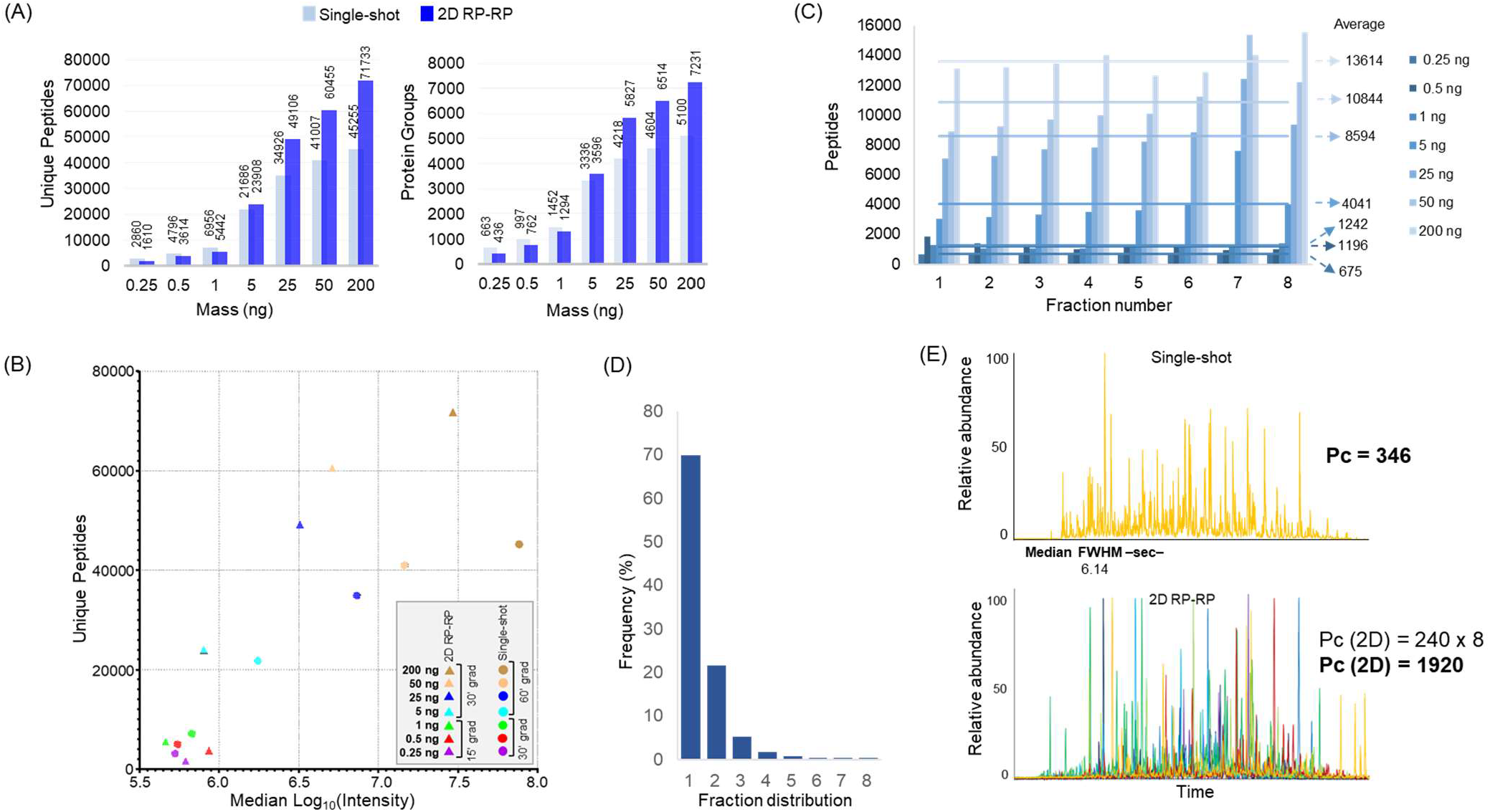
Performance of the 2D RP-RP at different human cell lysate digest sample amounts. (A) Number of unique peptides and protein groups identified by single-shot and 2D RP-RP applied to 0.25-200 ng sample amounts. (B) Number of unique peptides plotted as a function of median peptide intensities in single-shot and 2D RP-RP experiments according to the peptide sample amount. (C) Number of identified peptides across fractions. Numbers on the right indicate the average number of peptides identified per fraction according to the injected mass. (D) Histogram showing the observed frequency of peptides detected in one or multiple fractions. (E) Base-peak chromatograms of the single-shot and 2D RP-RP showing the median values of FWHM and the theoretical peak capacity (Pc) values of single-shot and 2D RP-RP.

### Performance at low sample amounts

The initial analysis suggested 5 ng (lower limit) and 50 ng of digest for sample loading represent appropriate choices for evaluating samples losses and performance of the 2D system with DDA analysis. We saw 25% and 26% increase in peptides and proteins for the 2D experiment over the single-shot experiment at 5 ng. For the 50 ng we identified 57,059 peptides and 6,598 proteins with the 2D experiment pipeline, corresponding to 62% and 30% increases in both peptides and proteins over single-shot (Figure 3A and Table S2). The vast majority of proteins and peptides observed in the single-shot experiment were also detected in the 2D experiment; with less than 5% being exclusive to the single-shot (Figure 3B). Generation of scatter plots of single-shot versus 2D RP-RP with the peptide and proteins identified with 5 ng and 50 ng of input material provided insights into performance. Interestingly, the intensity ratio between 2D and single-shot for both 5 ng and 50 ng at the peptide level had a value of 0.72 suggesting that, on average, the fractionation experiment had only lost 38% of the peptide signal (Figure 3C). The peptides observed exclusively in the single-shot experiment, as expected, correspond to those that have low intensities (Figure S5A). Conversely, the intensity distribution of the peptides observed exclusively by fractionation are low abundant as well (Figure S5B). Clearly, stochastic sampling continues to play a role in data acquisition. Reassuringly, we observe that with both 5 ng and 50 ng of input material, an increase in reported collective intensity for proteins with 2D compared to single-shot, most likely due to the increased number of peptides identified per protein, which confirms the value of fractionation (Figure 3D). Considering we opted to analyse the same amount by both approaches it was not surprising the observed dynamic range does not change significantly when switching to fractionation. It is noteworthy that the 2D fractionation system is capable of extending the experimental dynamic range by increasing sample loading, while with single-shot increasing loading quickly reaches saturation. The data suggests that the reason of increased identifications by fractionation is complexity reduction, at least for the sample amounts tested. In the single-shot runs there are many more components competing for ionisation and the charge available per scan (dictated by the AGC) is split across more peptides, reducing signal-to-noise for peak detection.

**Figure 3.**
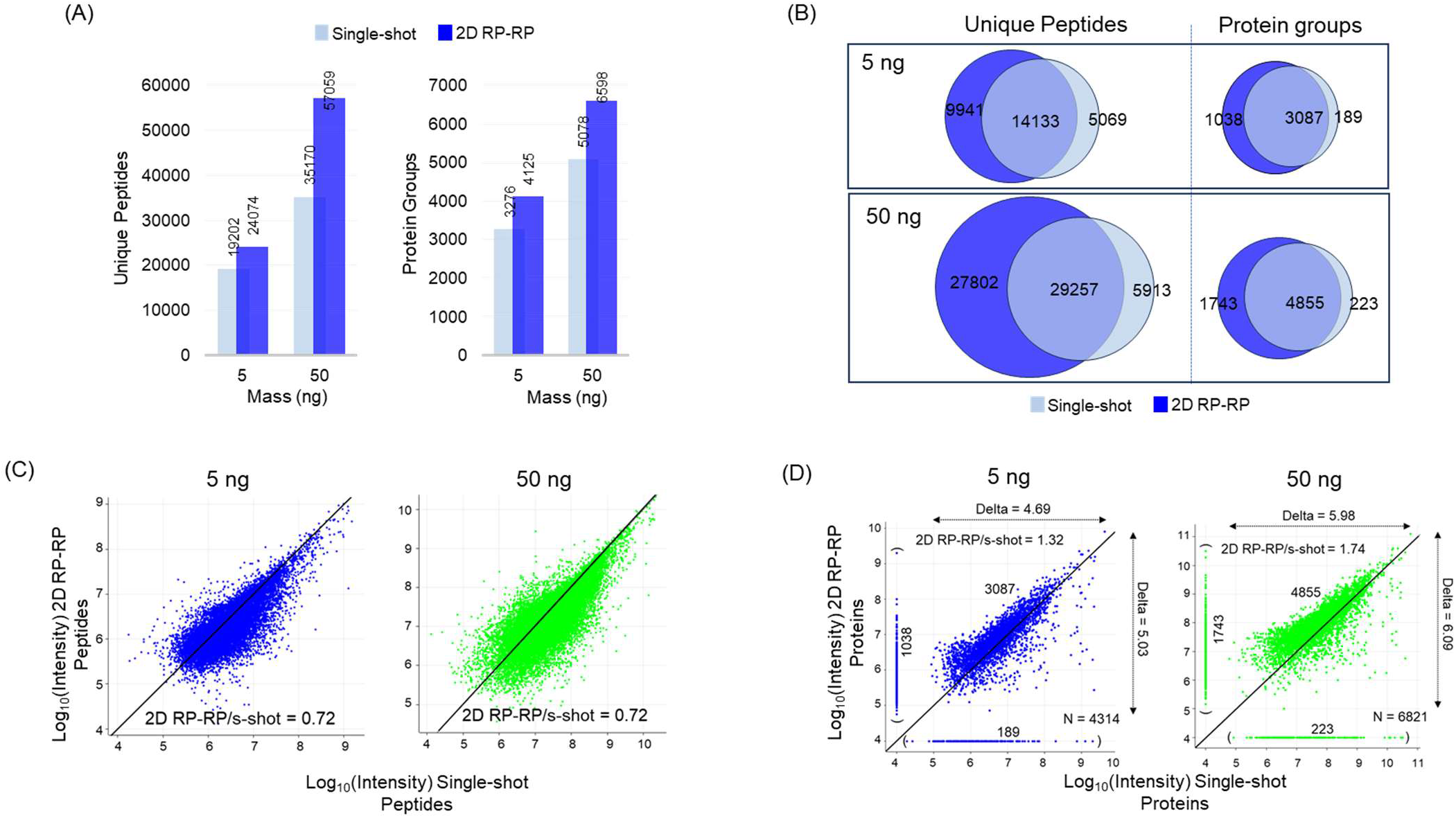
Performance of the 2D RP-RP in the fractionation of 5 and 50 ng of human cell lysate digested peptides. (A) Number of unique peptides and protein groups identified by single-shot and 2D RP-RP. (B) Venn diagrams showing the overlap of identified unique peptides and proteins between single-shot and 2D experiments. (C) Scatter plots of the peptides quantified in single-shot and 2D experiments showing the average intensity ratio between single-shot and 2D. (D) Scatter plots of the proteins quantified in single-shot and 2D experiments. On the X axis (bottom, in brackets) the graphs show the number of proteins only quantified by single-shot, while on the Y axis (left, in brackets) the number of proteins only quantified by 2D. The horizontal and vertical double headed arrows show the dynamic range of the single-shot and 2D experiments.

### Use of DIA and WWA combined with 2D RP-RP

The intensity scatterplots suggest that the advantage of 2D fractionation is the increase in proteome coverage due to the reduction in total signal splitting. Removing the need to select peaks for fragmentation as required by DDA approaches and increasing the AGC for the MS2 might alleviate the precursor selection issue of DDA as shown countless times by the DIA community.^39^ Thus, we repeated the 5 ng and 50 ng experiments but now using DIA and wide window acquisition approaches (WWA). See Table S2 for the list of identified peptides using DIA and WWA. The 5 ng experiments were similar in performance to DDA in our hands, with approximately 20,000 peptides and 3,500 proteins identified. However, 2D RP-RP combined with DDA appeared to be distinctly superior (Figure 4A). Strikingly, both DIA and WWA were substantially better than DDA for the 50ng single-shot experiment; both approaches nearly matched the performance of the 2DDDA experiment. The 2D RP-RP experiments for both DIA and WWA showed an increase of approximately 15% at the peptide level and protein level compared to the single-shot. Considering it gets progressively more difficult to increase the number of proteins observed as deeper the analysis becomes, such a gain is significant. We also calculated ‘completeness’ of the identified peptides for both DDA and WWA and observed an increase in missing values for peptide intensity with WWA. Over 50% of identified peptides did not have a corresponding intensity value for the single-shot experiment while that dropped to 38% for 2D. (Figure S6). Comparing the peptides observed across all acquisition approaches, approximately 53% of data is common among all approaches (Figure 4B). Interestingly, similar uniqueness levels between single-shot and 2D are also observed for DIA and WWA to the above DDA experiments (Fig 4C). In our hands WWA performed similar to DIA^40, 41^ suggesting both approaches are compelling choices as alternatives to DDA even when considering 2D RP-RP experiments.

**Figure 4.**
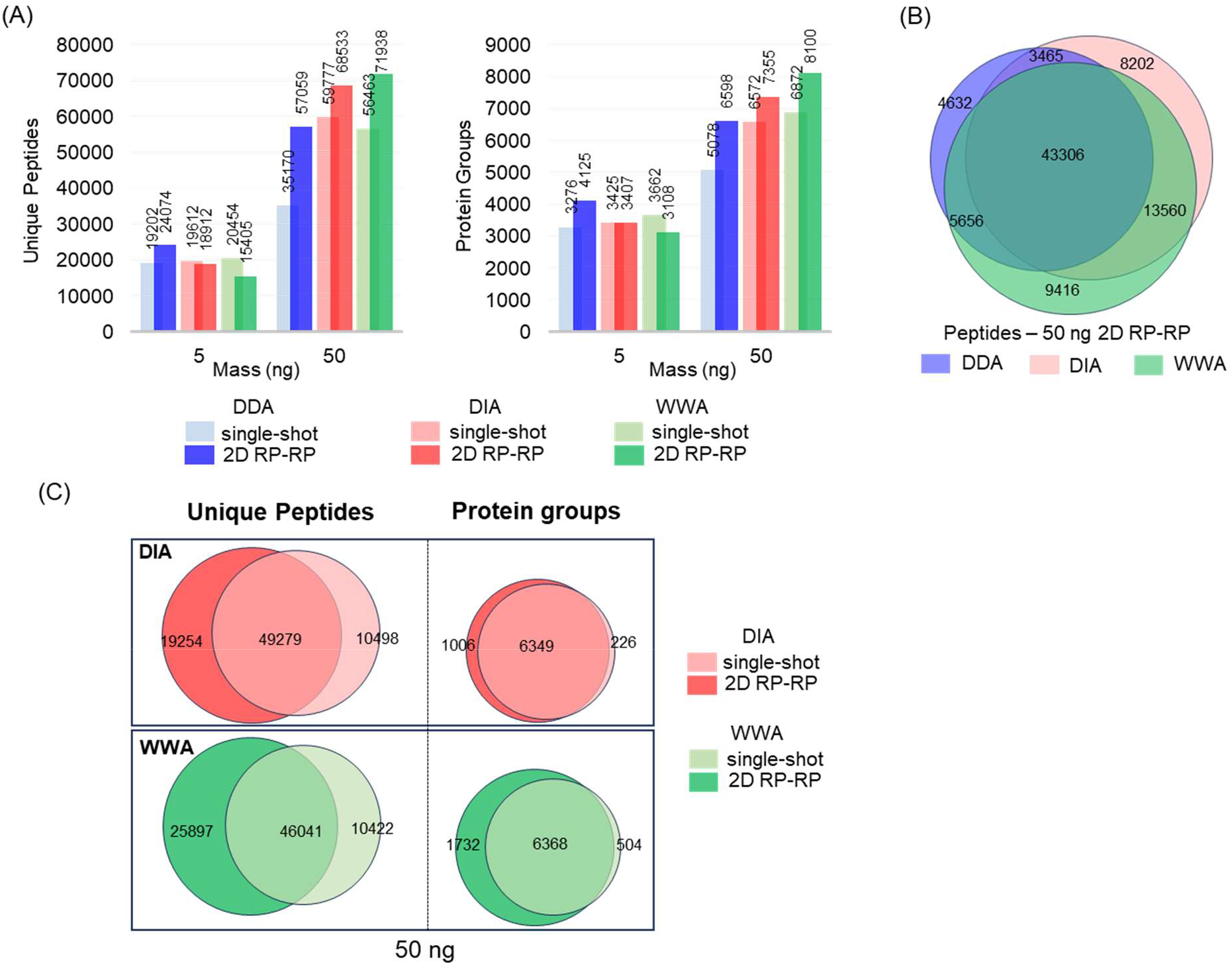
Performance of DIA and WWA in combination with 2D RP-RP. (A) Number of unique peptides and protein groups identified by single-shot and 2D RP-RP from 5 and 50 ng of Human Cell lysate digest samples. (B) Venn diagram showing the overlap of identified peptides across DDA, DIA and WWA from the fractionation of 50 ng of starting material. (C) Venn diagrams showing the overlap of identified unique peptides and proteins between 50 ng single-shot and 2D experiments using DIA and WWA approaches.

### Investigation of the use of enriched samples and AP-MS with the fractionation system

PTM enrichment is another potential class of experiments that would benefit from a low loss fractionation system. We performed a Zr-IMAC enrichment^24^ on a lysate to isolate the phosphoproteome and subjected 50 μg starting material to a single-shot (60-minute gradient) and 2D experiment (30-minute gradient, 8 fractions, total 4h). PTM type experiments disproportionately benefits more from increased sequencing coverage as sites are a peptide-centric property not protein-centric and as expected we saw significant increase in number of phosphosites observed in the 2D RP-RP experiment. Single-shot identified 9,929 sites and the 2D experiment identified 21,207 sites (Figure 5A and Table S3). Overlap between single-shot and 2D suggested loss of a significant number of sites (2,140 phosphosites) (Figure 5B). The exclusive sites observed for single-shot and 2D are in the lower intensity range; nevertheless, the 2D approach substantially identifies more new unique sites than those that are lost (Figure S7).

**Figure 5.**
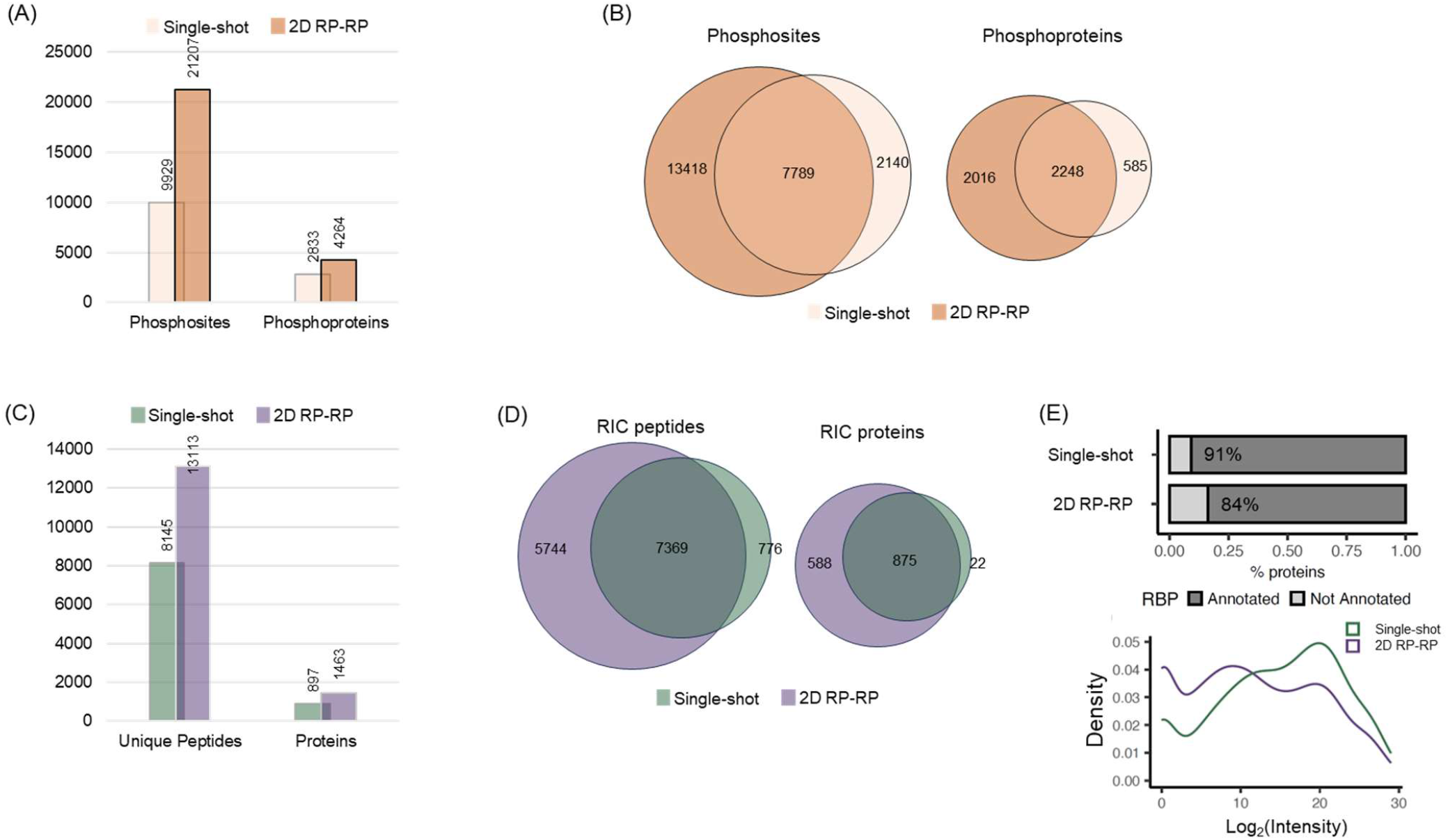
In-depth analysis of subproteomes by 2D RP-RP. (A) Number of phosphosites and phosphoproteins identified by single-shot and 2D experiments in Zr-IMAC enriched HeLa phosphopeptide sample. (B) Venn diagrams showing the overlap of identified phosphosites and phosphoproteins between single-shot and 2D. (C) Number of unique peptides and proteins identified by single-shot and 2D RP-RP from the analysis of HEK293 RNA-binding proteins isolated by RNA interactome capture technique. (D) Venn diagrams showing the overlap of identified peptides and RNA-binding proteins between single-shot and 2D. (E) Percentage of annotated RBPs in the set of proteins identified by single-shot and 2D (top), and the intensity distributions of RBPs identified by single-shot and 2D (bottom).

We also tested for a protein enrichment experiment by applying 2D system to an RNA interactome capture (RIC)^25, 42^ experiment, which enriches for RNA-binding proteins in cultured cells (HEK293 cells in this case). RIC employs irradiation of cell monolayers with ultraviolet (UV) light to promote “zero distance” RNA-to-protein crosslinks, followed by capture of protein-RNA complexes using oligo(dT) magnetic beads and elution by heat and RNases. Single-shot proteomic analysis revealed 897 RBPs. Strikingly, RBP identification increased to 1,463 when our 2D approach was applied, with over 98% of RBPs identified in the single-shot being also identified in the fractionated samples. Over five hundred additional RBPs were identified in the 2D fractionated run over the single-shot experiment, suggesting a substantial increase in depth (Figure 5C, D and Table S3). More than 84% of proteins identified in the fractionated run have been previously identified as RBPs (defined as occurring in 3 or more datasets in RBPbase^25^), which is slightly lower than with the single-shot analysis (91%, Figure 5E, top). The lower hit rate, therefore, suggests cautious approach to be taken for assigning RBP status and validated by orthogonal methods. Alternatively, it indicates that 2D may enable the identification of low abundance or sub-stoichiometric RBPs that conventional single-shot approaches miss. Supporting this, 2D fractionation increased coverage of proteins classified as RBPs that are expressed at low levels in cells (Figure 5E, bottom). This might reflect the identification of low abundant RBPs that are often missed in the absence of sufficient depth. Altogether, these results highlight the power of our 2D fractionation method to improve our understanding of the cellular proteome.

## CONCLUSIONS

We have developed a versatile and cost-effective 2D RP-RP system that enables deep proteome coverage from low amounts of material. In combination with DDA LC-MS analysis, the use of our fractionation system led to a substantial increase in peak capacity and identification rates over single-shot experiments for the analysis of human cell lysate, demonstrating superior proteome coverage using only 5 ng of digest with performance increasing with increasing amounts of material. We envision that our 2D setup can be readily applied to various DDA centric proteomics workflows and could be particularly useful for isobaric multiplexed experiments (e.g. TMT) applied to small cell populations, where sample availability is the issue. The application of DIA and WWA to fractionated samples also suggested value in the use of fractionation with the 2D WWA experiment allowing identification of more than 8,000 proteins from just 50 ng of material. Furthermore, the low loss 2D system demonstrably benefits phosphoproteomics and pull-down experiments, expanding coverage of both subproteomes relative to single-shot approaches.

## ASSOCIATED CONTENT

### Supporting Information

The Supporting Information is available free of charge on the ACS Publications website.

## METHOD

Sample preparation, Phosphopeptide enrichment, RNA Interactome Capture, Off-line High-pH reversed-phase nano-fractionation system, Nano-Liquid Chromatography and Mass Spectrometry, Data Analysis.; **FIGURE S1:** Evosep One HP pump pressure profile using a commercial EV-1106 and an in-house packed column at high and low-pH LC-MS (30SPD). Overlap of the base peak chromatograms corresponding to the high-pH LC-MS/MS analyses of Expi293F digest.; **FIGURE S2:** Column stability under basic conditions. Retention time alignments of five peptides identified in Expi293F digest by high-pH LC-MS/MS.; **FIGURE S3:** Orthogonality plot showing the retention times of Expi293F peptides analysed by high and low-pH LC-MS/MS using 30SPD method.; **FIGURE S4:** Nano-fractionation system overview, showing the Evosep One connected to the 3D-printer-based fraction collector; 3D-printed column bracket and platform.; **FIGURE S5:** Histograms of the intensities of peptides identified by single-shot and 2D RP-RP LC-MS/MS DDA analyses of 5 and 50 ng of Expi293F digest.; **FIGURE S6:** Number of peptides identified by single-shot and 2D RP-RP using DDA and WWA approaches showing the numbers of valid and missing values.; **FIGURE S7:** Histograms of the intensities of phosphosites identified by single-shot and 2D RP-RP LC-MS/MS DDA analyses from the Zr-IMAC enriched HeLa phosphopeptide sample.

## AUTHOR INFORMATION

### Author contributions

E.S.K. and S.M.: Conceptualization; G.N., E.S.K., and S.M. designed the fraction collector system; G.N. converted the 3D printer into a fraction collection system and created a dedicated GitHub page to document the system; E.S.K., Y.D., K.M.K. and L.I. performed the experiments; E.S.K. and Y.D. collected the data; S.C. provided technical support; E.S.K., L.I. and S.M. analysed the data; E.S.K. and S.M. wrote the initial draft; All authors read, edited and approved the final version of the paper.

### Notes

The authors declare no competing financial interest(s).

## ACKNOWLEDGMENTS

S.M, E.S.K, Y.D, S.C. and K.M.K were supported by EPSRC (V011359/1 (P)). A.C. is funded by the European Research Council (ERC) Consolidator Grant ‘vRNP-capture’ N# 101001634, the Career Development Award #MR/L019434/1, the John Fell Funds from the University of Oxford and the MRC grants MR/R021562/1 and MC_UU_00034/2.

