## Supporting Information for "Cheap, Robust and Versatile Two-Dimensional Chromatography system for Proteomics of Nanogram Scale Samples"

### Table of Contents

**METHOD:** Sample preparation, Phosphopeptide enrichment, RNA Interactome Capture, Off-line High-pH reversed-phase nano-fractionation system, Nano-Liquid Chromatography and Mass Spectrometry, Data Analysis.

**FIGURE S1:** Evosep One HP pump pressure profile using a commercial EV-1106 and an in-house packed column at high and low-pH LC-MS (30SPD). Overlap of the base peak chromatograms corresponding to the high-pH LC-MS/MS analyses of Expi293F digest.

**FIGURE S2:** Column stability under basic conditions. Retention time alignments of five peptides identified in Expi293F digest by high-pH LC-MS/MS.

**FIGURE S3:** Orthogonality plot showing the retention times of Expi293F peptides analysed by high and low-pH LC-MS/MS using 30SPD method.

**FIGURE S4:** Nano-fractionation system overview, showing the Evosep One connected to the 3D-printer-based fraction collector; 3D-printed column bracket and platform.

**FIGURE S5:** Histograms of the intensities of peptides identified by single-shot and 2D RP-RP LC-MS/MS DDA analyses of 5 and 50 ng of Expi293F digest.

**FIGURE S6:** Number of peptides identified by single-shot and 2D RP-RP using DDA and WWA approaches showing the numbers of valid and missing values.

**FIGURE S7:** Histograms of the intensities of phosphosites identified by single-shot and 2D RP-RP LC-MS/MS DDA analyses from the Zr-IMAC enriched HeLa phosphopeptide sample.

### METHOD

**Sample preparation and protein digestion.** Expi293F cells (Gibco) were re-suspended in a lysis buffer composed of 8M urea, 100mM ammonium bicarbonate and Roche Protease inhibitor cocktail and sonicated in Bioruptor Ultrasonicator (Bioruptor Pico, Diagenode) at 4°C using ultra-low sonication frequency for 30 cycles of 30 second sonication. After centrifugation at 14,000 x g, 4°C for 15 minutes, the supernatant was collected, and the protein concentration was estimated by BCA assay (Pierce). Proteins were reduced with 10 mM tris(2-carboxyethyl)phosphine (TCEP) for 30 minutes at room temperature followed by alkylation with 50 mM 2-chloroacetamide for 30 minutes in the dark. The urea concentration was diluted to 4M with 100 mM ammonium bicarbonate and the first step of digestion was carried out with LysC protease (Wako Pure Chemical Corporation, 125-05061) for 2 hours at 37°C with a protein to enzyme ratio of 50:1 (w/w). Samples were then further diluted to 1.3M urea (final concentration) and digested with trypsin (MS grade; Promega, cat. no. V5280) for 4 hours at 37 °C with a protein to enzyme ratio of 50:1 (w/w). Peptide samples were cooled to 4 °C and acidified with formic acid (FA) to a final concentration of 5% (v/v) and centrifuged at 16,200 × g for 10 minutes to remove any precipitate.

HeLa cells were re-suspended in a lysis buffer composed of 50mM Tris-HCl, pH 7.4, 150mM NaCl, 1% NP-40, 0.25% Triton X-100, 1mM DTT supplemented with protease and phosphatase inhibitors cocktails (P8340 and P0044, Merck). After sonication and protein quantification, 200 µg of proteins were reduced and alkylated as previously described, followed by SP3 method[1], using magnetic carboxylate-modified beads (GE45152105050250 and GE65152105050250, mixed at 1:1 ratio; Merck) with a protein to bead ratio of 4:1 (w/w). After cleanup, samples were re-suspended in 50 mM ammonium bicarbonate buffer and digested at 37°C for 4 hours using trypsin and LysC with a protein to enzyme ratio of 40:1 (w/w). Peptide digests were collected, and the beads were rinsed with 2% DMSO. Peptide samples were acidified with FA to a final concentration of 1% (v/v) and centrifuged at 16,200 × g for 10 minutes.

**Phosphopeptide enrichment.** Peptide sample from the digestion of HeLa cell lysate was submitted to Zr-IMAC phosphopeptide enrichment as previously described.[2] Briefly, peptides were desalted on HLB cartridge (Waters) and eluted with loading buffer (80% acetonitrile (ACN), 6% trifluoroacetic acid (TFA), and 1M glycolic acid) at a final concentration of 0.45µg/µL. Twenty microliters of Zr-IMAC beads (MR-ZHP005, MagReSyn) were equilibrated three times with 200µL of loading buffer, then added to the peptide mixture and incubated for 20 minutes in a ThermoMixer (Eppendorf), room temperature, at 1,350 rpm. Subsequently, beads were washed once with 400µL of loading buffer for 2 minutes, twice with 400µL of wash buffer 1 (80% ACN, 1% TFA) and twice with 400µL of wash buffer 2 (10% ACN, 0.2% TFA). Finally, peptides were eluted three times, each time by incubating beads with 80µL of elution buffer (4% ammonia hydroxide) for 10 minutes at room temperature, at 1,350 rpm. Eluate was acidified with FA and desalted on HLB cartridge as previously, except for elution where 25µL of 50% ACN solution was used.

**RNA Interactome Capture and sample preparation.** RNA-binding proteins (RBPs) present in HEK293 cells were profiled via RNA interactome capture (RIC) approach.[3, 4] HEK293 cells (2x$10^7$) were grown in 2x15cm dishes and incubated for 24 hours in 2% FBS DMEM. RIC samples were treated with benzonase for 30 minutes at room temperature to degrade both RNA and DNA and then processed via SP3 clean-up method followed by on-bead trypsin/LysC digestion as described before. Processed peptides were acidified by FA (final concentration 5%) prior to downstream analysis.

**Off-line High-pH reversed-phase nano-fractionation system.** Peptide samples re-suspended in 5% FA/0.015% n-Dodecyl-β-D-maltoside (DDM) solution were loaded on Evotip pure according to the manufacturer's instructions, and then separated by high-pH chromatography using an Evosep One system (Evosep Biosystems) and analysed at 30SPD method (44-minute gradient) using a double fritted in-house packed C-18 Reprosil-Gold column (150 µm ID x 150mm L, 1.9 µm particle size, Dr. Maisch). Mobile phases A and B consisted of 10 mM of triethylammonium bicarbonate (TEAB, pH8), and 100% ACN, respectively. Eluted peptides were fractionated into a 96-well plate using a 3D-printer-based fraction collector, adapted from a Creality Ender 5 S1 printer and powered by a Raspberry pi computer. The software is written in Python and uses PyGame for the GUI, Numpy of mathematical operations and PySerial for communication over USB. The list of components and instructions to build the fractionator can be found on github https://github.com/garethnisbet/Fraction-Collection-Unit. The 96-well plates were preloaded with 47.5µl of a solution composed by 5% FA/5% DMSO. Forty fractions were collected and automatically concatenated by the fraction collector into 8 main fractions at 1-minute intervals by combining fractions 1, 9, 17, 25, 33; 2, 10, 18, 26, 34; and so on.

**Nano-Liquid Chromatography, Mass Spectrometry and Data Processing.** Peptides were separated on Ultimate 3000 RSLCnano system (Thermo Fisher Scientific) equipped with a C-18 PepMap100 trap column (300 µm ID x 5mm L, 100Å, Thermo Fisher Scientific) and an in-house packed Reprosil-Gold C-18 analytical column (50 µm ID x 500mm L, 1.9 µm particle size, Dr. Maisch). Mobile phases (A: 0.1% FA, 5% DMSO and 94.9% water; B: 0.1% FA, 5% DMSO, 94.9% ACN) were delivered at a flow rate of 100 nL/min. For single-shot LC-MS/MS analysis, 30-minute gradient (10-36% B) was applied to peptide amounts ≤ 1ng and 60-minute gradient (10-33% B) was applied to amounts ≥ 5ng. For fractionated samples, peptides were separated using 15- (12-40% B) or 30-minute gradients for peptide amounts ≤ 1ng and ≥ 5ng, respectively. Eluting peptides were electro sprayed into an Orbitrap Ascend Tribrid mass spectrometer (Thermo Fisher Scientific), using Data Dependent (DDA), Data Independent (DIA) or Wide Window Acquisition (WWA) modes.

For DDA mode, full MS scans (350-1,400 m/z) were acquired in the Orbitrap analyzer at 60,000 resolution with a 1.2 x $10^6$ AGC target, and 123 ms maximum injection time. The twenty (15-minute gradient), thirty (30-minute gradient) or forty (60-minute gradient) most intense precursors (charge states 2-7) from MS1 scans were selected and isolated at 1.2 Th with the quadrupole for MS/MS event using higher-energy collision dissociation (HCD) at a normalized collision energy (NCE) of 26%, and the fragmentation spectra were detected by ion trap using turbo scan rate mode (2 x $10^4$ AGC target (1 x $10^4$, 15-minute gradient) and 32 ms maximum injection time). Dynamic exclusion was enabled with the following settings: exclusion duration = 15 s (15-minute gradient), 20 s (30- and 60-minute gradient), mass tolerance = ± 10 ppm, repeat count = 1.

In the case of WWA, we used the same acquisition parameters of DDA, except the ddMSn scan, where the isolation window was adjusted to 8 Th, using HCD fragmentation followed by orbitrap detection at 15,000 resolution.

For DIA measurements, full MS scans (400−1000 m/z) were acquired at 60,000 resolution with a 4 x $10^5$ AGC target, and 100 ms maximum injection time. Precursors were isolated with an isolation width of 8 m/z or 12 m/z for 60- and 30-minute gradients, respectively, using 63 or 42 windows covering a mass range of 400−900 m/z. Precursors were fragmented by HCD at a NCE of 28%. MS2 scans were acquired by Orbitrap at resolution of 15,000 with an AGC target of 7.5 x $10^5$ and maximum injection time of 27 ms.

Raw files from LC-MS/MS analysis using DDA mode were processed by FragPipe computational platform (version 20.0) with MSFragger[5, 6] (version 3.8) against the Uniprot human proteome reference database (Proteome ID: UP000005640) downloaded in August 2022 (79,759 sequences). Common contaminant proteins and an equal number of reversed sequence decoys were appended using Philosopher (version 5.0.0).[7] N-terminal acetylation and methionine oxidation were set as variable modifications and cysteine carbamidomethylation as fixed modification. Enzyme specificity was set to stricttrypsin (trypsin/P), precursor and fragment mass tolerance of 20 ppm with mass calibration and parameter optimization enabled, up to one missed cleavage, and minimum peptide length of 7 amino acids and maximum of 3 variable modifications were allowed per peptide. Other parameters remained at the default settings. False discovery rates were estimated and filtered for <1% at the peptide and protein levels. Raw files originating from the analysis of phosphopeptides-enriched samples were processed using MaxQuant software[8] (version 2.3.0.0) under default parameters. False-discovery rates were controlled at 1% both on peptide spectral match (PSM) and protein levels. Peptides with a minimum length of seven amino acids were considered for the search. Phospho(STY), N-terminal acetylation, methionine oxidation and were set as variable modifications and cysteine carbamidomethylation as fixed modification. Enzyme specificity was set to trypsin/P. A maximum of two missed cleavages were allowed.

DIA raw files were analyzed by DIA-NN software[9] (version 1.8.1) using library-free search. Spectral library was predicted in silico from the same Uniprot human proteome reference database used in DDA searches, covering the mass range between 300 to 900 m/z. For direct comparison with DDA searches, all remaining parameters, such as enzyme specificity, peptide modifications, charge states, and peptide length, were kept identical to the DDA workflow. FDR were estimated and filtered for <1% at the precursor and protein levels.

WWA raw files were processed by INFERYS[10] rescoring and CHIMERYS, implemented in Proteome Discoverer 3.0 software (Thermo Fisher Scientific), using the same protein database and search parameters employed in both DDA and DIA searches.

Raw files from this study have been deposited to ProteomeXchange Consortium via the PRIDE[11] partner repository with the data set identifier PXD051148. Subsequent analysis of data was performed in the Perseus environment[12] (version 1.6.15.0) and GraphpadPrism (version 9.3.1).

### Supplemental Figures

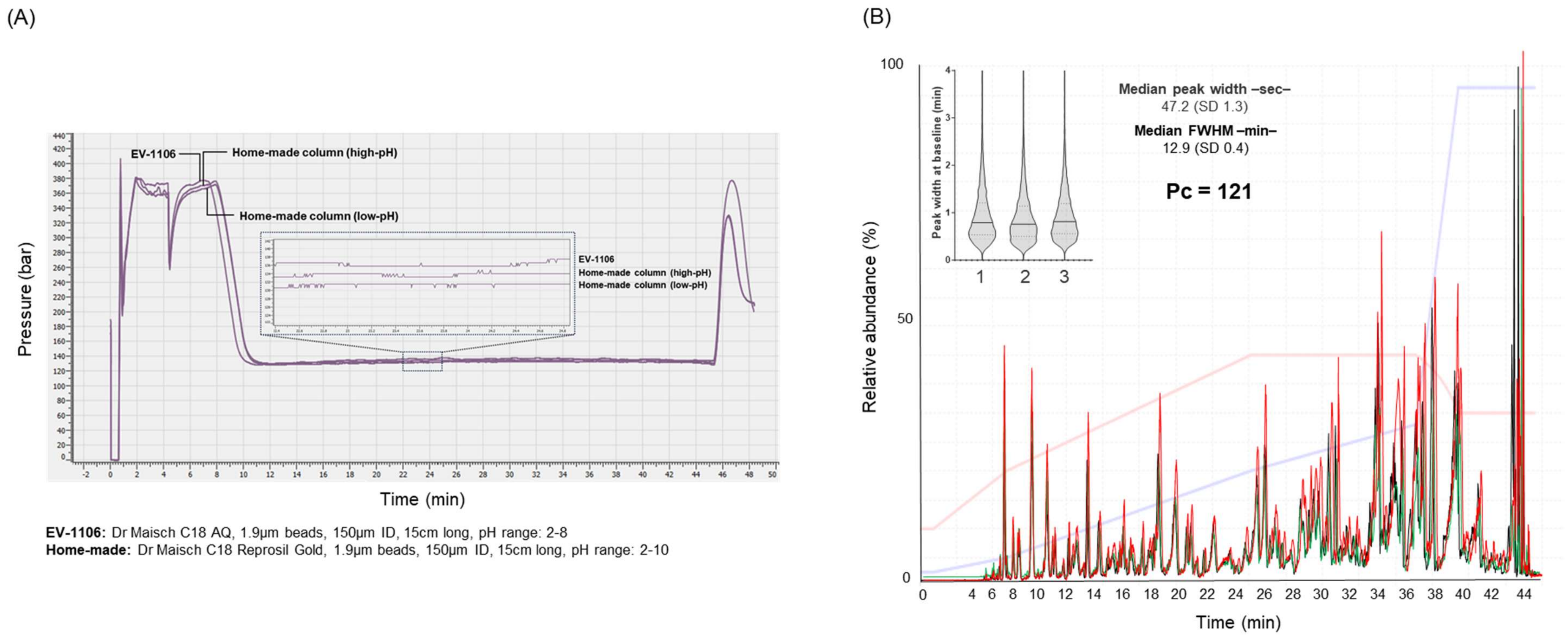


**Figure S1.** (A) HP pump pressure profiles for the separations performed by a commercial (EV-1106), and an in-house packed column at low and high-pH. (B) High-pH base peak chromatograms from three independent LC-MS/MS of Expi293F digests (50ng) using a 44-minute gradient (30 SPD) in Evosep One LC system. Peak capacity (Pc) value was calculated according to Kovalchuk et al. Mol Cell Proteomics. 2019 Feb;18(2):383-390.[13]

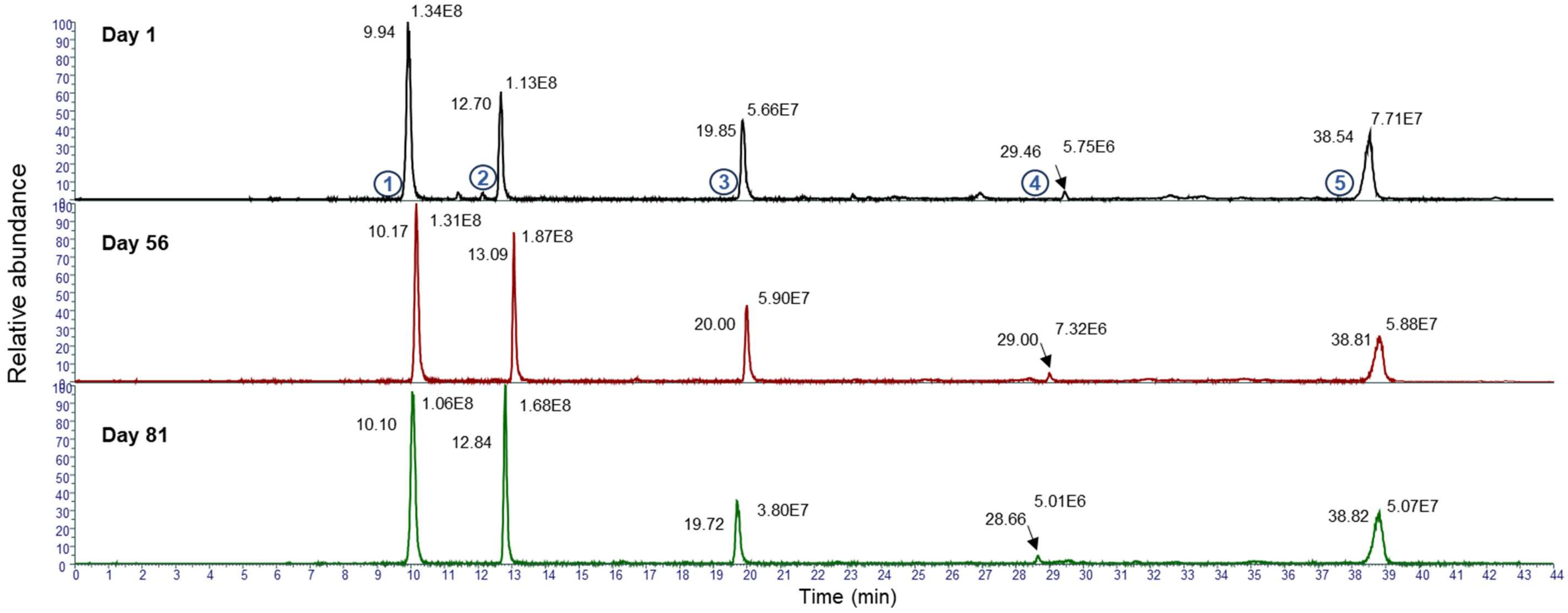


| Peak | m/z | Charge | Peptide | Average Intensity | Av RT (min) |
|---|---|---|---|---|---|
| 1 | 902.483 | 1 | STLEPVEK | 1.2E+08 | 10.06 |
| 2 | 1675.73 | 1 | ATAGDTHLGGEDFDNR | 1.6E+08 | 12.87 |
| 3 | 940.462 | 1 | DISLSDYK | 5.1E+07 | 19.86 |
| 4 | 1146.6 | 1 | DIDEVSSLLR | 6.0E+06 | 29.03 |
| 5 | 1614.81 | 1 | AFYPEEISSMVLTK | 6.2E+07 | 38.72 |

**Figure S2**. Column stability under basic conditions. Retention times and peak intensities of five peptides detected in Expi293F digest, monitored over 81 days, by high-pH LC-MS analysis (30SPD method).

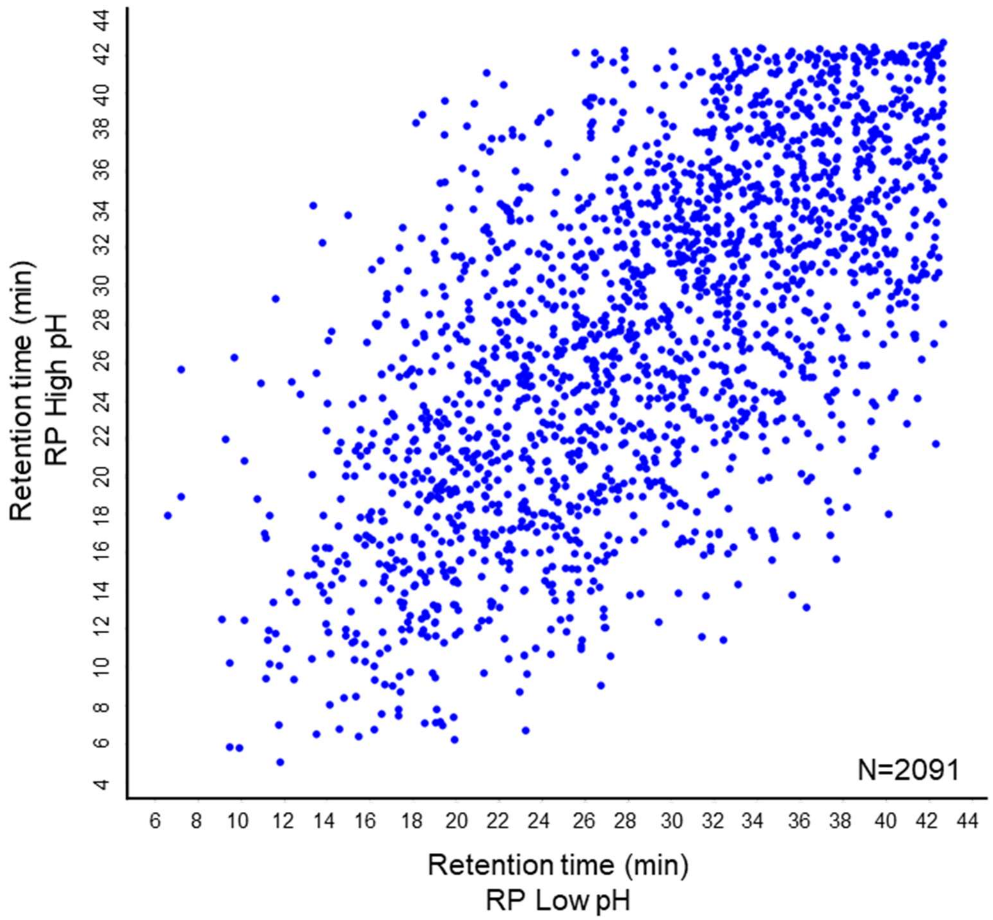


**Figure S3**. Orthogonality plot showing the retention times of Expi293F peptides separated by high and low-pH reversed phase chromatography using 30SPD method.

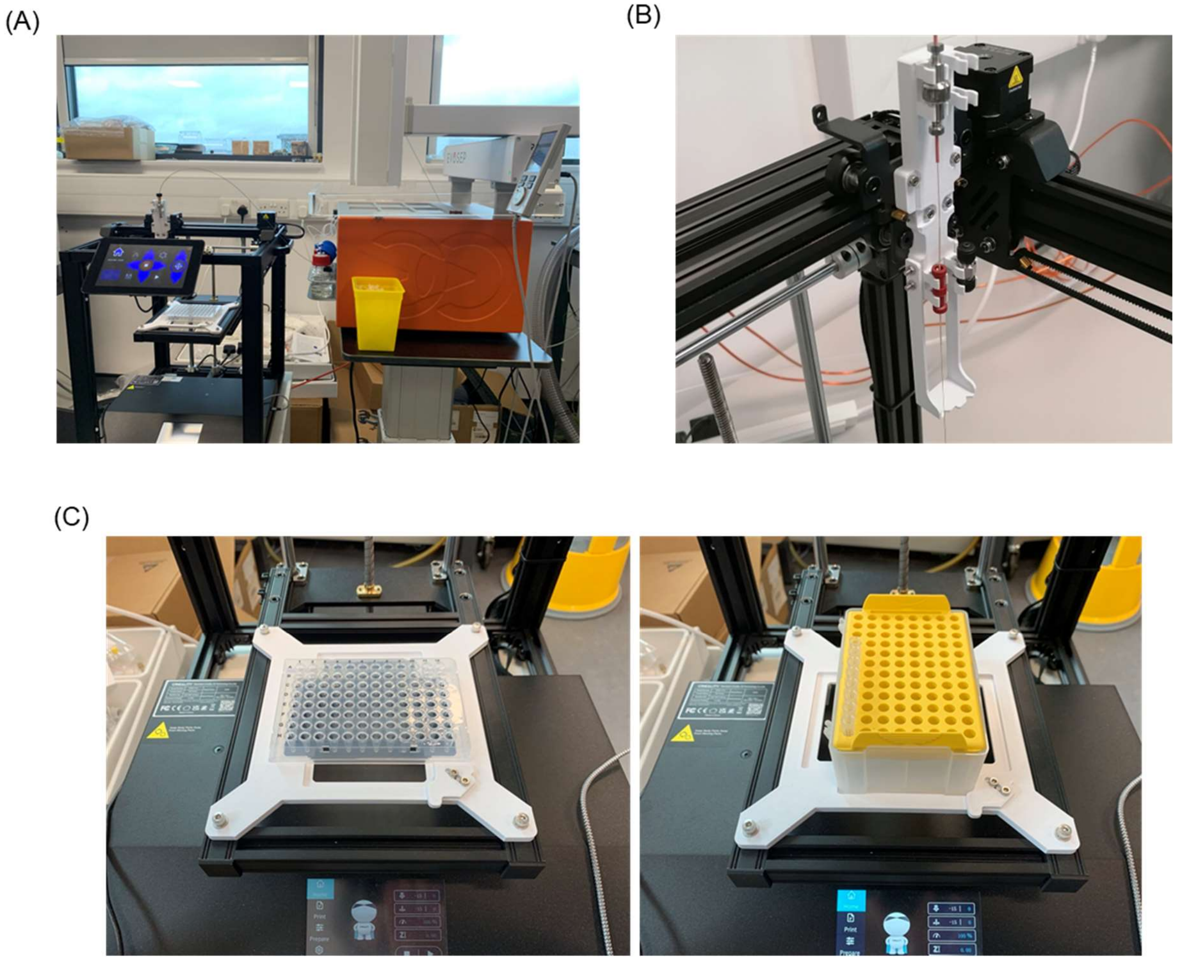


**Figure S4.** (A) Overview of the nano-fractionation system, consisting of the Evosep One and a modified 3D printer (Creality Ender 5 S1) fraction collector. (B) 3D-printed column bracket attached to the dismantled printer head to support the microcapillary column. (C) 3D-printed platform accommodates 96-well plates (left) or 96-Evotip racks (right).

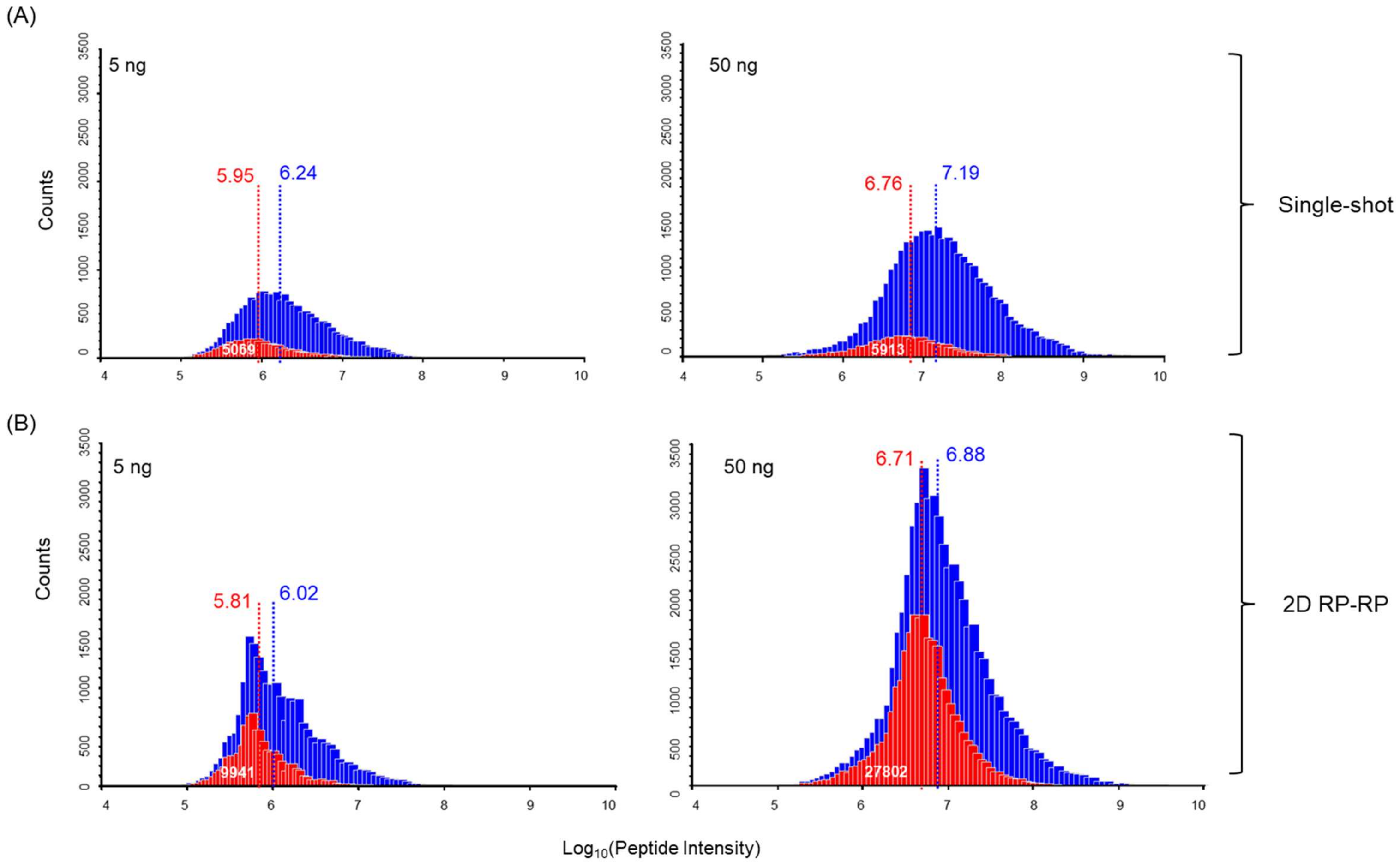


**Figure S5**. Intensity distribution of the peptides identified by single-shot (A) and 2D RP-RP (B) LC-MS/MS DDA analyses of 5 and 50 ng of Expi293F digest. The blue histograms show the intensity distribution of all peptides identified by single-shot or 2D RP-RP, while the red histograms show the distribution of intensities for peptides observed exclusively by each approach. Numbers on the top of the histograms indicate the median $Log_{10}$ intensities, while the number of exclusively identified peptides is shown in white.

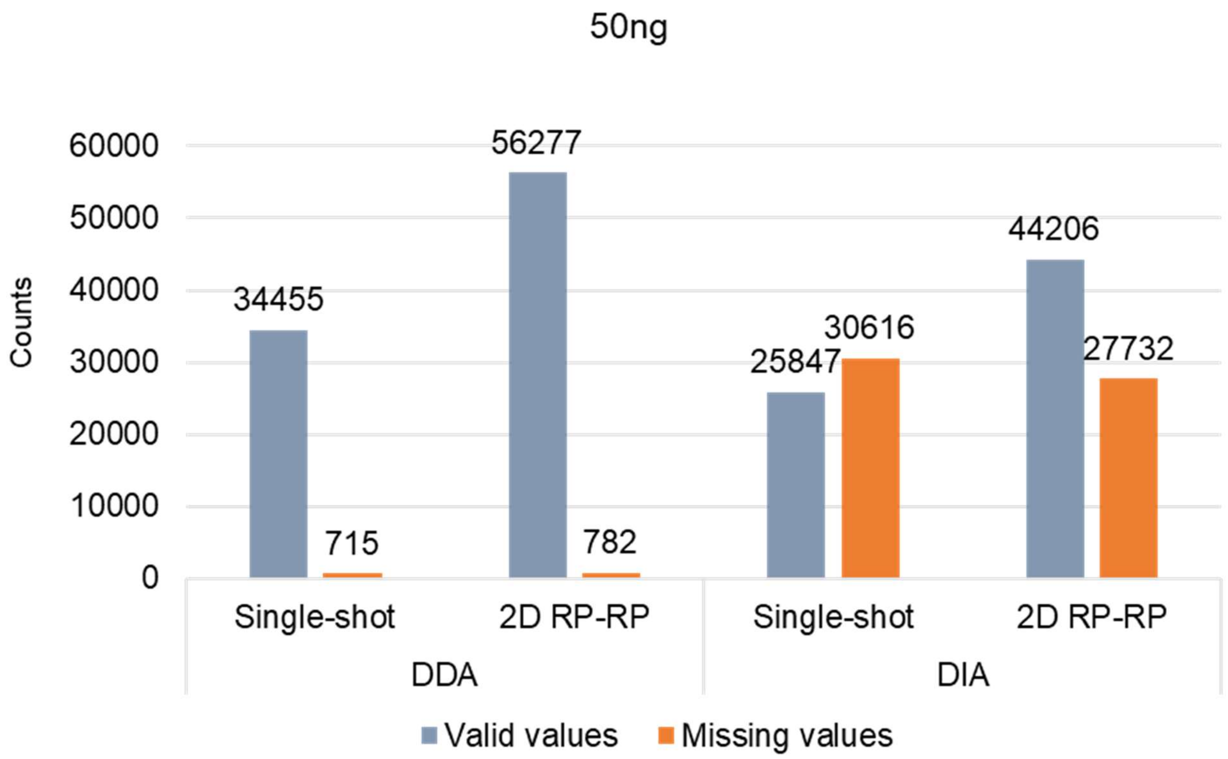


**Figure S6**. Number of peptides identified by the LC-MS/MS analysis of 50ng of Expi293F digest in single-shot and 2D RP-RP using DDA and WWA approaches showing the numbers of valid and missing values for peptide intensity.

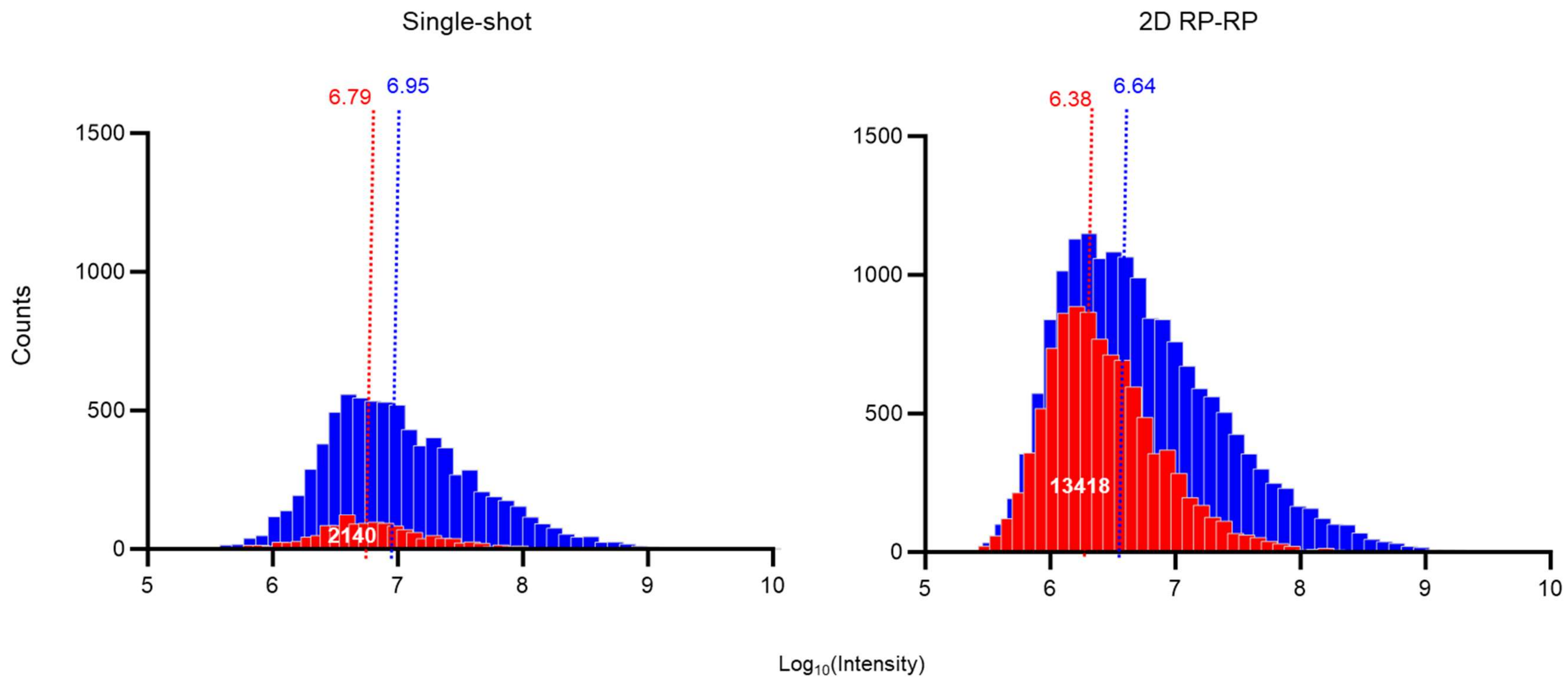


**Figure S7.** Intensity distribution of the phosphosites identified by single-shot (left) and 2D RP-RP (right) LC-MS analyses of Zr-IMAC enriched HeLa phosphopeptide sample. The blue histograms show the intensity distribution of the whole set of phosphosites identified by single-shot or 2D RP-RP, and in red, the distribution of the sites observed exclusively by each approach. Numbers on the top correspond to the median $Log_{10}$ intensity values and in white, the number of unique phosphosites in single-shot and 2D RP-RP.
